# N-terminal protein acetylation by NatB modulates the levels of Nmnats, the NAD^+^ biosynthetic enzymes in *Saccharomyces cerevisiae*

**DOI:** 10.1101/814327

**Authors:** Trevor Croft, Padmaja Venkatakrishnan, Christol James Theoga Raj, Benjamin Groth, Timothy Cater, Su-Ju Lin

## Abstract

NAD^+^ is an essential metabolite participating in cellular biochemical processes and signaling. The regulation and interconnection among multiple NAD^+^ biosynthesis pathways are not completely understood. We previously identified the N-terminal (Nt) protein acetyltransferase complex NatB as a NAD^+^ homeostasis factor. Cells lacking NatB show an approximate 50% reduction in the NAD^+^ level and aberrant metabolism of NAD^+^ precursors, which are associated with a decrease of nicotinamide mononucleotide adenylyltransferases (Nmnat) protein levels. Here we show this decrease in NAD^+^ and Nmnat protein levels is specifically due to the absence of Nt-acetylation of Nmnat (Nma1 and Nma2) proteins, and not other NatB substrates. Nt-acetylation is a critical regulator of protein degradation by the N-end rule pathways, indicating absence of Nt-acetylation may alter Nmnat protein stability. Interestingly, the rate of protein turnover (t_1/2_) of non-Nt-acetylated Nmnats does not significantly differ from Nt-acetylated Nmnats, suggesting reduced Nmnat levels in NmatB mutants are not due to increased post-translational degradation of non-Nt-acetylated Nmnats. In line with these observations, deletion or depletion of N-rule pathway ubiquitin E3 ligases in NatB mutants is not sufficient to restore NAD^+^ levels. Moreover, the status of Nt-acetylation does not alter the rate of translation initiation of Nmnats. Collectively our studies suggest absence of Nt-acetylation may increase co-translational degradation of nascent Nmnat polypeptides, which results in reduced Nmnat levels in NatB mutants. Nmnat activities are essential for all routes of NAD^+^ biosynthesis. Understanding the regulation of Nmnat protein homeostasis will facilitate our understanding of the molecular basis and regulation of NAD^+^ metabolism.

## INTRODUCTION

NAD^+^ and its reduced form NADH play an essential role in coupling oxidation to energy producing metabolism. NADP^+^ and NADPH are derivatives of NAD^+^(H) and are important for the biosynthesis of large molecules and anti-oxidant defense. Additionally, NAD^+^ is consumed as a substrate in NAD^+^-dependent reactions. Some of these reactions, like sirtuin-mediated deacetylations, have signaling functions by modification of their protein substrates [1, 2]. Therefore, NAD^+^ level can modulate cell signaling through these NAD^+^-dependent reactions. Cells must properly balance NAD^+^ biosynthesis with its consumption in order to maintain optimal cellular function. Moreover, abnormalities in NAD^+^ metabolism have been associated with a number of metabolic disorders and disease [3–6]. Understanding the regulation of NAD^+^ homeostasis may help elucidate the mechanisms of these disorders.

NAD^+^ biosynthesis in yeast is maintained by three major pathways: *de novo* biosynthesis, nicotinic acid (NA) and nicotinamide (NAM) salvage [7, 8] and nicotinamide riboside (NR) salvage [9] (Fig. 1*A*). NAD^+^ biosynthesis by the *de novo* pathway begins at tryptophan and requires Bna proteins to synthesize nicotinic acid mononucleotide (NaMN). This pathway is the most resource intensive of the three and is heavily regulated [10–12]. NaMN is also produced by the NA/NAM salvage branch, which begins with precursors like NA and NAM. Under NA abundant conditions, NA/NAM salvage is the preferred NAD^+^ biosynthesis route, and *BNA* genes are silenced by the NAD^+^-dependent deacetylase Hst1 [10, 11]. NaMN produced from both branches is converted to nicotinic acid adenine dinucleotide (NaAD) by nicotinamide mononucleotide adenylyltransferases (Nmnats) Nma1 and Nma2 [13–16], followed by the amidation to NAD^+^ by Qns1 [17, 18]. In the NR salvage pathway, NR is phosphorylated by NR kinase Nrk1 to produce nicotinamide mononucleotide (NMN) [9]. Nma1, Nma2, or Pof1 transfer the AMP moiety of ATP to NMN to produce NAD^+^ [13–16, 19]. It has been shown that Nma1 and Nma2 have dual substrate specificity towards both NMN and NaMN [13–15] whereas Pof1 activity is specific for NMN [19]. NR can also enter the NAM salvage branch by nucleosidases Uhr1 and Pnp1 [20]. Moreover, yeast cells also release and re-uptake small NAD^+^ precursors such as NA, NAM, QA and NR (Fig. 1*A*) [11, 19, 21]. Specific transporters Tna1 (for NA and QA) [22, 23] and Nrt1 (for NR) [24] are responsible for the uptake of NAD^+^ precursors whereas the mechanisms of precursor release remain unclear. Together three branches of NAD^+^ biosynthesis pathways are coordinated and provide the cell with NAD^+^ tuned to the cellular requirements and environmental conditions.

**FIGURE 1.**
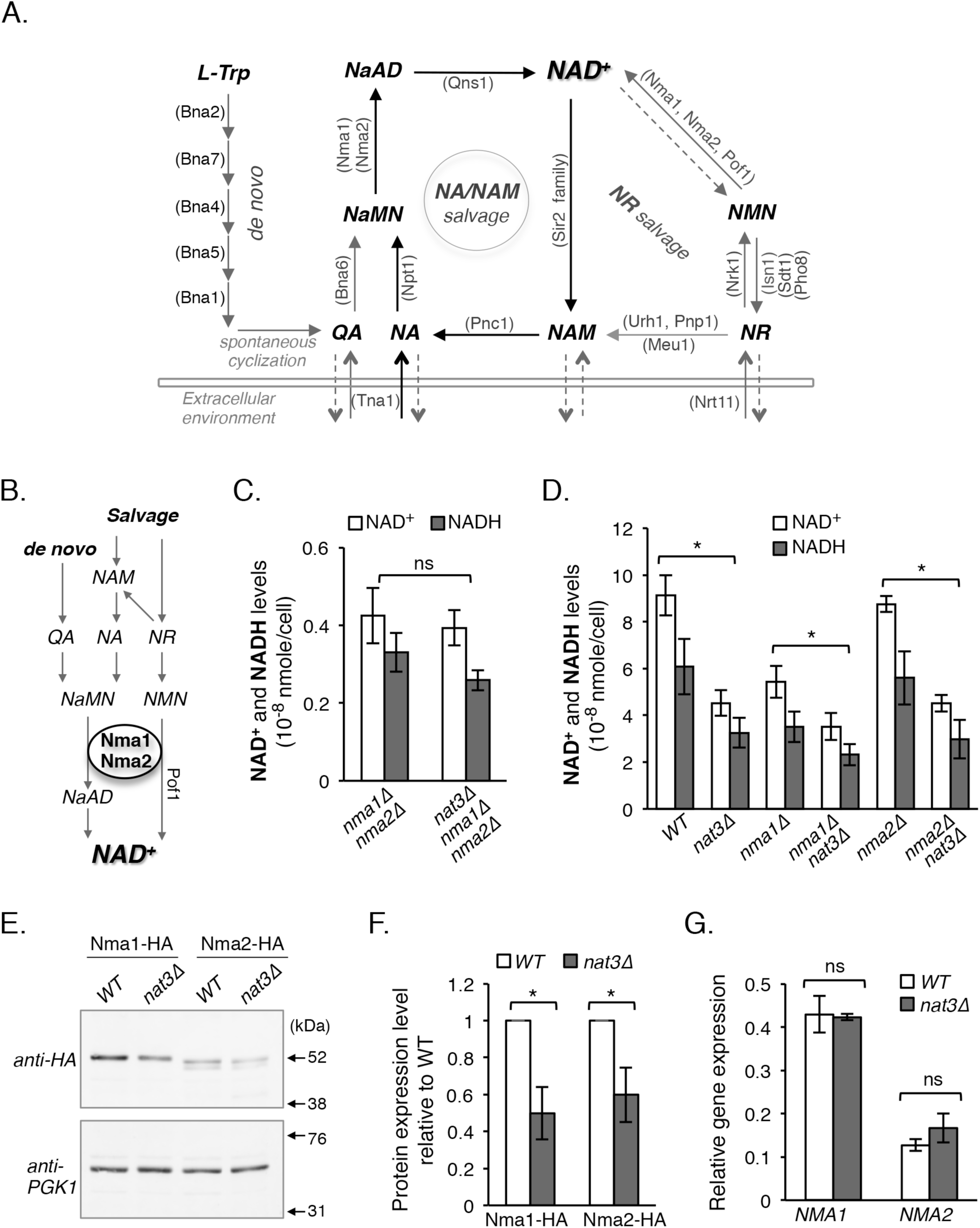
The NAD^+^ deficiency of the NatB mutant is primarily due to Nma1 and Nma2 defects. *A*, a simplified model of NAD^+^ biosynthesis pathways in *Saccharomyces cerevisiae*. NAD^+^ can be synthesized *de novo* from tryptophan (*L-Trp*), and by salvaging NAD^+^ intermediates through the NA/NAM and NR cycles. Yeast cells also release and re-uptake small NAD^+^ precursors, however the mechanisms are not completely understood. NA, nicotinic acid. NAM, nicotinamide. QA, quinolinic acid. NR, nicotinamide riboside. NaMN, nicotinic acid mononucleotide. *NaAD*, deamido NAD^+^. *NMN*, nicotinamide mononucleotide. Abbreviations of protein names are shown in *parentheses*. *B*, a schematic diagram showing that all 3 NAD^+^ biosynthesis pathways require Nma1 and Nma2 activities. A third Nmnat Pof1 is part of the NR salvage pathway exclusively. *C*, *nma1Δnma2Δ* and *nma1Δnma2Δnat3Δ* mutants have similar levels of NAD^+^(H). NR was supplemented to the cell culture at 10 µM. Cells lacking *NMA1* and *NMA2* can grow in the presence of NR using Pof1. *D*, NAD^+^(H) levels of *nma1Δ* and *nma2Δ* were compared combining *nma1Δ* or *nma2Δ* with *nat3Δ* deletions. Both *nma1Δnat3Δ* and *nma2Δnat3Δ* had lower levels of NAD^+^ in comparison with the single Nmnat deletion. *E*, Western blot analysis of Nma1 and Nma2 tagged with HA in combination with PGK1 control in both WT and *nat3Δ* cells. Arrows mark the positions of the molecular weight markers. *F*, Quantitative analysis of *E* showing Nma1 and Nma2 protein depletions due to *nat3Δ* deletion. Nma1-HA and Nma2-HA are normalized to PGK1 and WT is set to 1. *G*, gene expression qPCR analysis of *NMA1* and *NMA2* mRNA in WT and *nat3Δ* cells. Data shown are representative of at least three independent experiments. The *error bars* denote standard deviations. The *p* values are calculated using Student’s *t* test (*, *p* < 0.05).

The Nmnat proteins Nma1 and Nma2 are the only enzymes that are responsible for generation of NAD^+^ in all three biosynthesis pathways (Fig. 1*B*). Therefore, Nma1 and Nma2 may be critical for the co-regulation of *de novo*, NA/NAM salvage and NR salvage pathways. In fact, we have previously shown that overexpression of *NMA1* caused a significant increase in the total NAD^+^ content of the cell [21]. It was also shown in a recent study that Nma1 is the only NAD^+^ biosynthesis enzyme that upon overexpression can increase the NAD^+^ content in the cell, and modulates NAD^+^ to correlate with the concentrations of ATP [12]. Overall, these studies suggest Nmnats are rate-limiting factors in NAD^+^ biosynthesis pathways. Interestingly, Nmnats appear to have additional roles in the cell. Overexpression of both *NMA1* and *NMA2* suppresses proteotoxicity in yeast models of polyglutamine and α-synuclein-induced neurodegeneration by targeting toxic misfolded proteins for degradation [25]. This supports the neuroprotective role that human Nmnats play in axon degeneration and neurodegenerative diseases [5, 26, 27].

Previously we identified the NatB complex, Nat3 (catalytic subunit) and Mdm20 (regulatory subunit), as a potential regulator of NAD^+^ biosynthesis, possibly through N-terminal (Nt-) acetylation of Nma1 and Nma2 [21]. The levels of Nma1 and Nma2 proteins were reduced in NatB deletion mutants compared to wild type (WT). Nt-acetylation is primarily a co-translational protein modification and carried out by a Nt-acetyltransferases [28]. Nt-acetylation neutralizes the positive charge carried by the free amino group making the N-terminus more hydrophobic. This protein modification has potential to alter folding, complex formation, localization, and degradation [28]. NatB carries out Nt-acetylation of approximately 15% of encoded yeast proteins, which are Met-retained peptides with Asp, Asn, Glu or Gln at position two. NatB mediated Nt-acetylation is efficient, and nearly all proteins that code as NatB substrates are indeed NatB substrates, which is not true for other Nt-acetyltransferases. Interestingly, yeast NatB substrates are predominantly, near 100%, found in the acetylated state [28, 29]. In line with these observations, we showed that Nma1 and Nma2 peptides identified from WT cells were 100% acetylated whereas Nma1 or Nma2 peptides identified from NatB mutants were 95%-100% non-acetylated [21]. Therefore, Nt-acetylation appears to be critical for maintaining proper Nmnat protein levels and thus NAD^+^ biosynthesis. To date, a direct link between NatB mediated Nt-acetylation of Nmnat and NAD^+^ biosynthesis has not been established since NatB also has many other substrates in the cell.

It also remains unclear why NatB mutants have lower Nmnat protein levels. It is possible non-Nt-acetylated Nmnats are more susceptible to degradation. N-terminal regions of proteins contain specific degrons, termed N-degrons [30]. These degrons are residue specific and are described by the N-end rule pathways. The Arg/N-end rule pathway is dedicated to peptides with non-Nt-acetylated N-terminal Arg, Lys, His, Leu, Trp, Phe, Ile, Tyr, or Met retained peptides followed by a bulk hydrophobic residue at position two. Ubr1 is the E3 ligase that targets these substrates for ubiquitination and subsequent degradation by proteasome. The Ac-N-end rule pathway is typically specific for Nt-acetylated substrates, which are targeted by the E3 ligases Doa10 or Not4. In absence of Nt-acetylation, Ac/N-end rule proteins may become unrecognizable by their E3 ligases and long-lived [31]. While Nma1 and Nma2 do not fulfill the requirements for degradation by the Arg/N-end rule pathway, it is worth noting that examples of Ac/N-substrates becoming Arg/N-substrates in the absence of Nt-acetylation have been observed [32, 33]. In addition to the possibility of being degraded by the N-end rule pathways, non-Nt-acetylated Nmnats may be degraded during translation since the absence of Nt-acetylation may affect its folding or complex formation. It has been shown that all newly synthesized polypeptides are monitored by a ribosome-based quality control system during translation [34–37]. As a result, significant portions of nascent polypeptides are subject to co-translational degradation [36, 38–40]. However detailed mechanisms of this protein quality control system are not completely understood.

Overall, these studies suggest that maintaining Nmnat protein levels is critical for proper NAD^+^ biosynthesis, and that Nt-acetylation maybe important for Nmnat protein stability. In this study, we aim to examine whether NatB regulates NAD^+^ biosynthesis directly though Nt-acetylaiton of Nma1 and Nma2 and why absence of Nt-acetylation causes the reduction of Nmnat protein levels in NatB mutants. Our studies help understand the roles of Nt-acetylation on Nmnats and provide a molecular basis for the regulation of NAD^+^ homeostasis factors.

## RESULTS

### Nma1 and Nma2 play a primary role in NatB deletion associated NAD^+^ deficiency

We previously reported altered NAD^+^ homeostasis in NatB deletion mutants, *nat3Δ* and *mdm20Δ* [21]. These NatB deletion mutants have low levels of NAD^+^ with an increased flux of NAD^+^ precursors through the vacuolar NR salvage branch, which promotes the formation and release of NAM and NA. This defect was associated with a decrease in Nma1 and Nma2 protein levels, both of which are NatB substrates [21] and catalyze a reaction shared by all three pathways (Fig. 1*B*). Nearly 15% of all yeast proteins are Nt-acetylated by NatB complex. To confirm that the low NAD^+^ phenotype in NatB deletion mutants is primarily due to Nma1 and Nma2 defects, we compared NAD^+^ levels of the *nma1Δnma2Δ* and *nma1Δnma2Δnat3Δ* mutants (Fig. 1*C*). If the low NAD^+^ levels in *nat3Δ* cells are due to additional defects, we would expect to see further decreases in NAD^+^ and NADH levels in the *nma1Δnma2Δnat3Δ* mutant. Under normal growth conditions the *nma1Δnma2Δ* mutants are inviable unless NR is supplemented to the growth media. It was shown that Pof1, a third Nmnat and not a NatB substrate, can convert NMN to NAD^+^ in the NR salvage branch [19]. Therefore, adding NR to growth media supports growth of the *nma1Δnma2Δ* mutants through the activity of Pof1 (Fig. 1*A*). In line with our expectations, there is no significant difference in NAD^+^ and NADH levels between the *nma1Δnma2Δ* and *nma1Δnma2Δnat3Δ* strains grown in NR supplemented media. Furthermore, to examine whether NatB activity is important for both Nma1 and Nma2 function, we paired the *nat3Δ* deletion with either *nma1Δ* or *nma2Δ* single deletion. As shown in Fig. 1*D*, both the *nma1Δnat3Δ* and *nma2Δnat3Δ* double mutants showed a near 50% decrease in NAD^+^ and NADH levels compared to the respective single Nmnat deletion alone (Fig. 1*D*). These results indicate that the low NAD^+^ defect in NatB mutants acts primarily through both Nma1 and Nma2.

Next we determined whether Nma1 and Nma2 protein levels were decreased to similar extent in NatB deletion *nat3Δ* mutant when a different control is employed. In previous work we used actin, whose expression appeared stable in NatB mutants, as loading control [21]. However, since actin is also a NatB substrate [42], we chose Pgk1 [41] for protein turnover studies in this work. Consistent with previous findings, Nma1 and Nma2 protein levels were reduced by nearly 50% when normalized to Pgk1 and WT (Fig. 1, *E* and *F*). Although Nt-acetylation is primarily a co-translational protein modification we also wanted to confirm that the low protein levels of Nma1 and Nma2 were not due to reduced transcription. As shown in Fig. 1*G*, no significant difference in the level of mRNAs (determined by RT-qPCR) was found when comparing *nat3Δ* mutant to WT. Together these results suggest that Nt-acetylation is required for maintaining NAD^+^ by modulating the Nma1 and Nma2 protein levels.

### NatB promotes NAD^+^ biosynthesis directly through Nt-acetylation of Nma1 and Nma2

We previously found overexpression of Nma1 can increase the levels of NAD^+^ in both WT and *nat3Δ* cells [21]. This result is supported by the finding that overexpression of Nma1 can bypass an ATP-dependent regulatory mechanism [12]. However, this may also indicate that the low NAD^+^ defect in NatB mutants is not directly due to lack of Nt-acetylation of Nmnats but may rather stem from lower intracellular concentrations of ATP or other unforeseen consequences since NatB also acetylates many other proteins. To test if the Nt-acetylation modification is indeed important for Nma1- and Nma2-mediated NAD^+^ biosynthesis, we replaced the endogenous Nma1 and Nma2 with mutant versions expected to be NatA substrates instead of NatB substrates (Fig. 2*A*). NatA substrates are not Met-retained like NatB substrates, but first undergo a processing by removal of the initiator methionine by methionine aminopeptidases, and the new amino-terminus is then acetylated by NatA (Fig. 2*A*, *left panel*) [28]. Additionally, potential NatB substrates (as predicted by their second amino acids) are almost always acetylated by NatB. This is not true for all potential NatA substrates. Based on reported proteomic data we chose to insert Ser between the first two amino acids, Met and Asp, in Nma1 and Nma2 (Nma1/2_MD_ ➔ Nma1/2_M<u>S</u>D_) (Fig. 2*A*, *right panel*) because this peptide sequence is almost always acetylated by NatA [28, 29].

**FIGURE 2.**
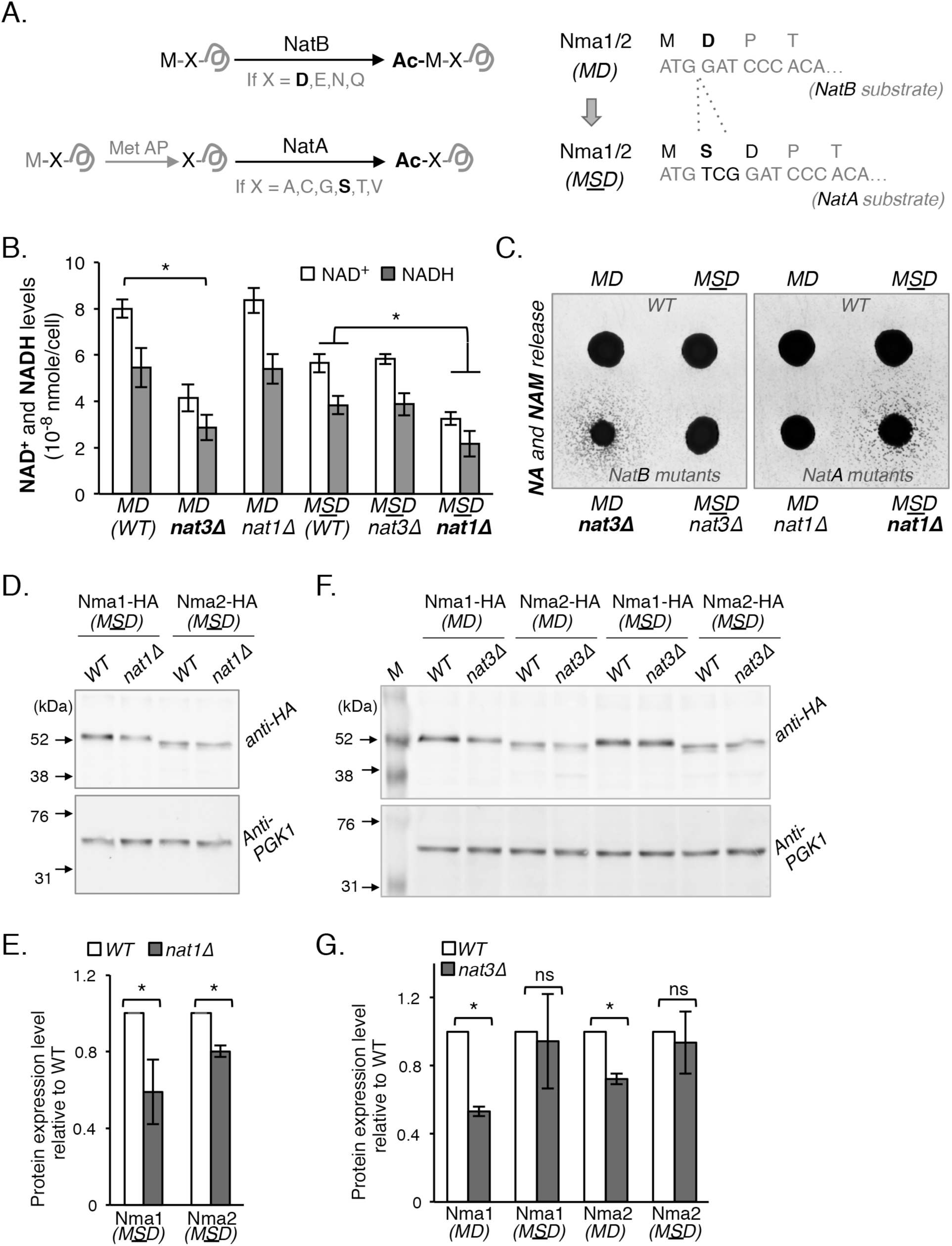
The NAD^+^ metabolism defect of the NatB mutant is directly linked to absence of Nt-acetylation and associated reduction of Nma1 and Nma2 protein levels. *A*, schematic showing of amino acid sequences that result in Nt-acetylation by NatB or NatA (*left panel*). NatB acetylates Met-retained peptides when the second amino acid is Asp (D), Asn (N), Glu (E), and Gln (Q). NatA acetylates substrates where the second amino acid is Ala (A), Cys (C), Gly (G), Ser (S), Thr (T), or Val (V). NatA substrates are first processed by methionine aminopeptidases MetAPs, which remove the N-terminal methionine. To make Nma1 and Nma2 become NatA substrates, Ser was inserted as the second amino acid (*right panel*), which converts Nma1/2_MD_ to Nma1/2_MSD_. *B*, NAD^+^(H) levels of *nma1Δnma2Δ cells* harboring NatB-specific (MD) and NatA-specific (MSD) Nma1 and Nma2 combined with NatB (*nat3Δ*) or NatA (*nat1Δ*) specific deletions. Changing Nma1 and Nma2 to NatA substrates makes NAD^+^(H) levels sensitive to *nat1Δ*, whereas *nat3Δ* no longer decreases NAD^+^(H). *C*, levels of NA/NAM release determined by a cell-based NA/NAM cross-feeding assay. NA/NAM-dependent recipient cells (*bna6Δnrk1Δnrt1Δ*) were plated as a lawn on niacin free SD. Next, *nma1Δnma2Δ* cells harboring NatB-specific (MD) or NatA-specific (MSD) Nma1 and Nma2 combined with *nat3Δ* or *nat1Δ* were spotted on top of the lawn as feeder cells. Growth of the recipient cells indicates the level of NA/NAM released by each specific feeder strains. *D*, levels of NatA-specific Nma1-HA_MSD_ and Nma2-HA_MSD_ are reduced in *nat1Δ* (a NatA mutant) cells determined by Western blot analysis*. E*, quantification of Western blot protein studies shown in *D*. NatA-specific Nma1-HA_MSD_ and Nma2-HA_MSD_ are normalized to PGK1 and WT is set to 1. *F*, levels of NatA-specific Nma1_MSD_ and Nma2_MSD_ are not significantly changed in *nat3Δ* cells (a NatB mutant). Western blot analysis of NatB-specific (MD) and NatA-specific (MSD) Nma1-HA and Nma2-HA in WT and *nat3Δ* cells. *G*, quantification of protein studies shown in *F*. protein levels are normalized to PGK1 and WT is set to 1. Data shown are representative of at least three independent experiments. The *error bars* denote standard deviations. The *p* values are calculated using Student’s *t* test (*, *p* < 0.05).

The *MD* (*WT*) strain carries Nma1 and Nma2 in their native form as NatB substrates, which is made by integrating pRG207-*NMA1_MD_ (WT)* and pRG205-*NMA2_MD_ (WT)* into *nma1Δnma2Δ* cells. The *MSD* (*WT*) strain carries modified Nma1 and Nma2 in a form of converted NatA substrates, and is made by integrating pRG207-*NMA1_MSD_* and pRG205-*NMA2_MSD_* into *nma1Δnma2Δ* cells. These strains were then paired with either *nat3Δ* (NatB mutant) or *nat1Δ* (NatA mutant) deletions for NAD^+^ measurements (Fig. 2*B*). As expected, the NAD^+^ levels of *MD* strains carrying *NMA1_MD_* and *NMA2_MD_*(NatB substrates) were decreased when combined with *nat3Δ* but not *nat1Δ* deletion, indicating Nma1_MD_ and NMA2_MD_ were still sensitive to deletions of *NAT3* but not *NAT1*. However, NAD^+^ levels of *MSD* strains carrying *NMA1_MSD_* and *NMA2_MSD_*(NatA substrates) were sensitive to deletions of *NAT1* but not *NAT3* (Fig. 2*B*). These results suggest the low NAD^+^ phenotype of NatB deletion mutants is due to the absence of Nt-acetylation on Nma1 and Nma2. Notably, the *MSD* (WT) strain has lower basal NAD^+^ levels compared to the *MD (WT)* strain, which suggests that acetylation of Nma1_MSD_ and Nma2_MSD_ by NatA is not optimal.

Next we examined whether observed NAD^+^ reduction in the *nat1Δ* mutant carrying *NMA1_MSD_* and *NMA2_MSD_*was due to a blockage in NA/NAM salvage. If so, this mutant is expected to show increased release of NA and NAM as previously reported for the *nat3Δ* mutant [21]. A cross-feeding reporter assay was employed to determine the level of NA and NAM released by cells. In this system, cell growth of the NA and NAM dependent mutant *bna6Δnpt1Δnrk1Δ* (recipient cells) was used as readout of the amount of NA and NAM released by the feeder cells (mutants of interests) [21]. In agreement with expectations, the growth of recipient cells was only observed when *nat3Δ* mutant harboring *NMA1_MD_* and *NMA2_MD_* (Fig. 2*C*, *left panel*) and *nat1Δ* mutant harboring *NMA1_MSD_* and *NMA2_MSD_* (Fig. 2*C*, *right panel*) were used as the feeders. Moreover, the *nat3Δ* mutant harboring *NMA1_MSD_* and *NMA2_MSD_*no longer releases more NA and NAM when compared with *nat3Δ* harboring *NMA1_MD_* and *NMA2_MD_* (Fig. 2*C*, *left panel*). Therefore it appears that NAD^+^ biosynthesis mediated by Nma1 and Nma2 requires Nt-acetylation by NatB for optimal product formation, and lack of this modification reduces the NAD^+^ content of the cell resulting in release of NAD^+^ precursors like NAM and NA into the environment.

To confirm if the protein levels of Nma1_MSD_ and Nma2_MSD_ were indeed depleted in the *nat1Δ* mutant, whole-cell lysates were generated from cells expressing c-terminally HA-tagged Nma1_MSD_ and NMA2_MSD_ followed by Western blot analysis. As shown in Fig. 2*D* and 2*E*, Nma1_MSD_ and Nma2_MSD_ proteins were depleted in *nat1Δ* cells to about 60% of the control strain. Importantly, Nma1_MSD_ and Nma2_MSD_ protein levels were not significantly depleted in *nat3Δ* mutants (Fig. 2, *F* and *G*). These results support a positive correlation between Nmnat protein levels and NAD^+^ levels. In addition, Nt-acetylation appears to play a direct role in maintaining the level of Nmnat proteins and NAD^+^ biosynthesis.

### Degradation by the N-end rule pathways is not the main cause of reduced Nma1 and Nma2 protein levels and NAD^+^ levels in NatB mutants

Next we examined whether Nma1 and Nma2 proteins are less stable and degraded at a faster rate in NatB mutants. To test if the N-end rule pathways play an important role in the reduction of Nma1 and Nma2 protein levels, E3 ligases associated with the Arg-N and Ac-N end rule pathways were deleted or depleted in combination with deletion of *NAT3*. If any of these E3 ligases were important for rapid ubiquitination and subsequent degradation of the non-acetylated Nma1 and Nma2 proteins, we expected to see NAD^+^ levels restored. As shown in Fig. 3*A*, deletions of E3 ligases *DOA10* (Ac-N) or *UBR1* (Arg-N) did not restore NAD^+^ levels. Next, since double deletions of *NOT4* (Ac-N) and *NAT3* were synthetically lethal, we employed an auxin-inducible degron (AID) system) [43, 44] to transiently deplete Not4 in *nat3Δ* mutant (Fig. *3B*). In this system, the minimum *AID* sequence [43, 44] was inserted to the chromosome region corresponding to the c-terminus of Not4 generating *NOT4-AID\** strain. Next, a plasmid carrying a specific E3 ubiquitin ligase *OsTIR1-9myc* was introduced to the *NOT4-AID\** strain. Upon addition, auxin (indole-3-acetic acid, IAA) binds to TIR and promotes the interaction of TIR and the AID degron of Not4 resulting in Not4 degradation. We first validated that Not4 was indeed depleted under our experimental conditions within 2 hours upon auxin induction by Western blot analysis (Fig. 3*B*, *upper right panel*). However, NAD^+^ levels were not restored in *nat3Δ* mutants by Not4 depletion (Fig. 3*B*). Therefore, it appears that Ubr1, Doa10 and Not4 may not play a major role in the rapid degradation of non-acetylated Nma1 and Nma2. Interestingly, a small but not significant increase of NAD^+^ was observed when Not4 was depleted in WT cells (Fig. 3*B*) suggesting the Ac-N rule pathway may play a role in maintain proper protein turnover of Nt-acetylated Nma1 and Nma2.

**FIGURE 3.**
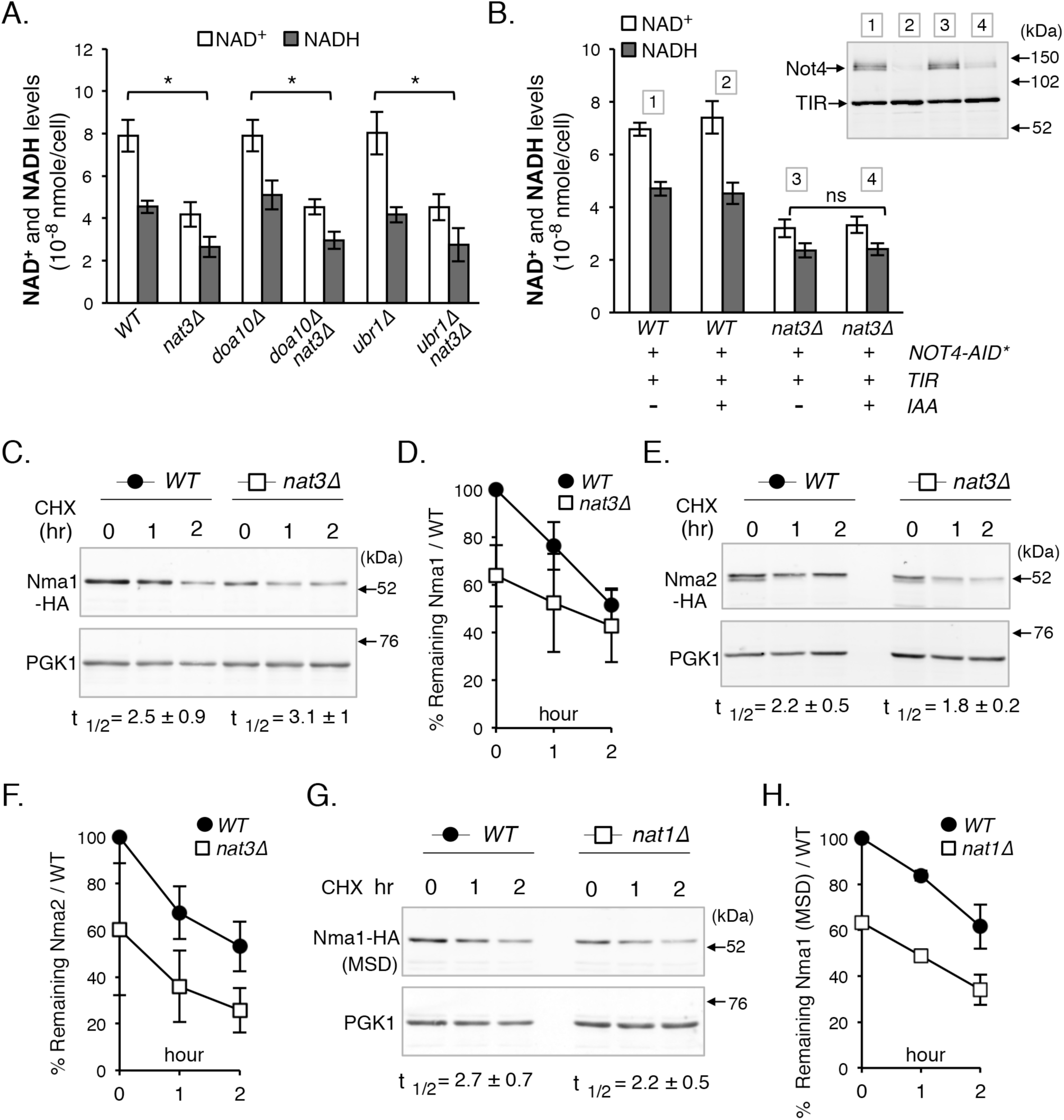
Post-translational degradation by the N-end rule pathways is not the main cause of reduced Nma1 and Nma2 proteins and NAD^+^ levels in NatB mutants. *A*, deletions of E3 ligases of the Ac/N-end rule pathway (*doa10Δ*) and Arg/N-end rule pathway (*ubr1Δ*) do not restore NAD^+^(H) levels in *nat3Δ* cells. *B*, depletion of Not4 (an Ac/N-rule E3 ligase) does not restore NAD^+^(H) levels in *nat3Δ* cells (*left panel*). Not4 depletion was induced in cells harboring Not4-AID*-myc9 and TIR by addition of auxin (IAA) to 1 mM for 2 hours (*upper right panel*). *C*, cycloheximide (CHX) chase studies of Nma1-HA in WT vs *nat3Δ* cells over 2 hours determined by Western blot. CHX was added to cell culture to a final concentration of 0.4 mg/ml. Half-life (t_½_) displayed is based on three independent experiments. *D*, quantification of protein studies shown in *C*. Protein levels are normalized to PGK1, and WT at 0 hour is set to 100%. *E*, cycloheximide chase studies of Nma2-HA in WT vs *nat3Δ* cells over 2 hours determined by Western blot. Half-life displayed is based on three independent experiments. *F*, quantification of protein studies shown in *E*. Proteins levels are normalized to PGK1, and WT at 0 hour is set to 100%. *G*, cycloheximide chase studies of NatA-specific Nma1-HA_MSD_. Half-life (t_½_) displayed is based on three independent experiments. *H*, quantification of protein studies shown in *G*. protein levels are normalized to PGK1, and WT at 0 hour is set to 100%. Data shown are representative of at least three independent experiments. The *error bars* denote standard deviations. The *p* values are calculated using Student’s *t* test (*, *p* < 0.05).

While no known E3 ligase of the two N-end rule pathways has a significant role in the rapid degradation of non-acetylated Nma1 and Nma2, we could not rule out the possibility of an unknown E3 ligase. To test the possibility that non-acetylated Nma1 and Nma2 were being targeted for rapid degradation compared to their acetylated counterparts, we employed cycloheximide chases followed by Western blot analysis to determine the half-lives (t_1/2_) of the proteins. A small but insignificant change in the rate of degradation was observed when comparing acetylated Nma1 and Nma2 with their non-acetylated counterparts (Fig. 3, *C* to *F*), which does not account for an approximate 50% reduction in Nma1 and Nma2 in NatB deletion mutants (Fig. 1, *E* and *F*). Additionally, we determined the half-lives of non-acetylated and acetylated Nma1_MSD_ in WT and *nat1Δ* cells (Fig. 3, *G* and *H*) and the results were in agreement with those shown in Fig. 3*C* to 3*F*. Therefore, it appears that the low levels of Nma1 and Nma2 proteins in NatB mutants are not due to more rapid post-translational protein degradation of the non-Nt-acetylated Nma1 and Nma2 by the N-rule pathways.

### Nt-acetylation may protect Nmnats from co-translational degradation

Overall our results suggest the non-Nt-acetylated counterparts of Nmnats may be rapidly degraded during translation. Supporting this, it has been shown that co-translational protein degradation is not related to the half-lives of mature folded proteins [36, 37], which is in line with our observation of Nmnats (Fig. 3, *C* to *H*). Moreover, it was shown that proteins with longer disordered regions at the N-terminus have a higher rate of co-translational degradation [36, 37], and that an intrinsically disordered region has been predicted at the N-terminus of Nma1 and Nma2 [45]. Therefore, it is possible that Nt-acetylation helps proper folding of Nmants during translation, and consequently, non-Nt-acetyalted Nmnats are more susceptible for co-translational degradation.

However, it is also possible that the NatB complex may facilitate Nmnat translation and therefore the levels of Nmnats are reduced in absence of NatB. In fact, Nt-acetyltransferase complexes have been shown to associate with non-translating and translating ribosome subunits [46]. To address this question, we determined the rate of translation of *NMA1* and *NMA2* mRNA by polysome profile analysis as described [47–49]. Cell lysate generated from WT and NatB mutant cells were fractionated on a 10%-50% sucrose gradient, which gave rise to the polysome profile shown in Fig. 4*A* (*WT*) and Fig. 4*B* *(nat3Δ)*. Sucrose gradient solutions were collected in 10 equal volume fractions; Fraction #1 corresponds to the top of the gradient (free mRNA) and fraction #10 corresponds to the bottom of the gradient. Fractions #7-10 correspond to translating mRNAs associated with heavy polysomes. The amount of gene-specific mRNA in each fraction was calculated to obtain the proportion (%) of the target mRNA found in each fraction as described [48, 50]. As a result, relative abundance and distribution of specific mRNA in 10 fractions were shown for *NMA1* (Fig. 4*C*), *NMA2* (Fig. 4*D*) and a non-NatB substrate control *UBC6* (Fig. 4*E*). Overall, the % of *NMA1* and *NMA2* mRNA associated with the polysome fractions (#7-10) were not significantly altered in the NatB mutant, indicating the Nt-acetylation status does not affect the rate of Nmnat translation. Interestingly, the % of *NMA1* and *NMA2* mRNA in the polysome fractions were higher than that of the *UBC6* control, suggesting both *NMA1* and *NMA2* are relatively actively translated. Together our characterization of Nma1 and Nma2 were in line with the properties of proteins that have higher rates of translation but are prone to co-translational degradation [36, 37]. Although the mechanisms of co-translational protein degradation are not completely understood, out studies of Nmnat suggest Nt-acetylation may play a role.

**FIGURE 4.**
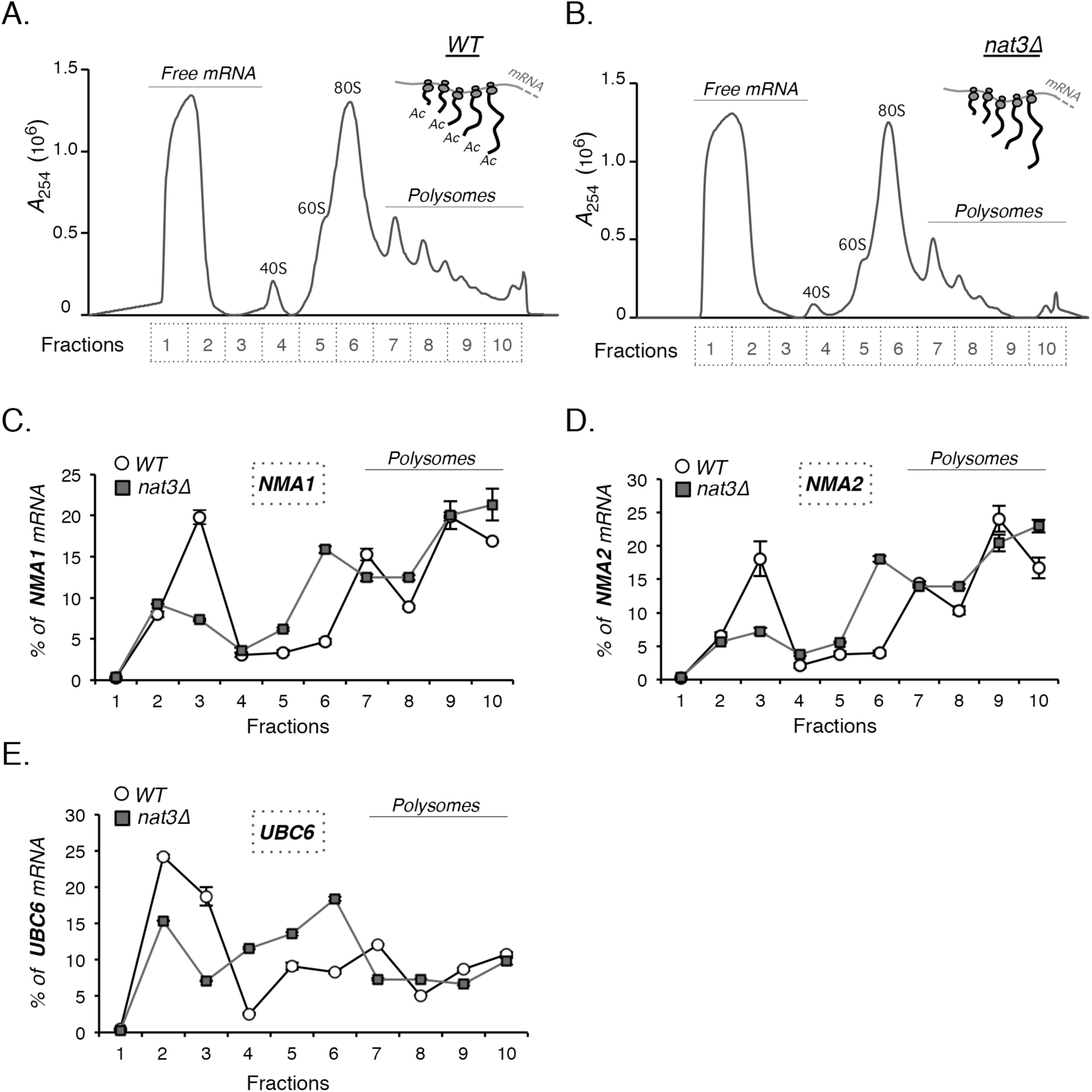
The status of Nt-acetylation does not alter the rate of translation initiation of Nmnats. *A*, polysome profile of WT cells. Optical density profile (*A*_254_) of WT cell lysate in 10% to 50% sucrose gradient is shown. Sucrose gradient solutions are collected in 10 equal volume fractions. Fraction #1 corresponds to the top of the gradient (free mRNA) and fraction #10 corresponds to the bottom of the gradient. Translated mRNAs are associated with the heavy polysome fractions (fractions # 7-10). 40S: small ribosomal subunit; 60S: large ribosomal subunit; 80S: ribosome (monosome). *B*, polysome profile of NatB mutant (*nat3Δ*) cells. Optical density profile (*A*_254_) of *nat3Δ* cell lysate in 10% to 50% sucrose gradient is shown. For *C*, *D* and *E*, mRNA coding for *NMA1* and *NMA2* are actively translated. *UBC6* is a control for non-NatB substrate. X axis indicates the number of fractions collected from the sucrose gradient. Y axis shows % of gene-specific mRNA present in each fraction. The sum of gene-specific mRNA in all 10 fractions is set at 100%. Data shown are representative of three independent experiments. The *error bars* denote standard deviations.

## DISCUSSION

In this study, we showed that Nt-acetyaltion by NatB is important for maintaining the levels of Nma1 and Nma2 proteins, and therefore NAD^+^ biosynthesis (Fig. 1). We also established a direct link between NatB and Nmnats in the regulation of NAD^+^ metabolism (Fig. 2). NatB Nt-acetylates ∼15% of cellular proteins, therefore observed NAD^+^ defects might be indirectly due to unknown off-target effects. In this regard, we asked whether we could alter NAD^+^ levels by modulating Nt-acetylation of Nma1 and Nma2 by replacing the endogenous Nma1 and Nma2 with mutant versions expected to be NatA substrates instead of NatB substrates. As expected, the NAD^+^ levels of strains carrying NatA-specific Nma1 and Nma2 (*MSD*) became sensitive to deletions of *NAT1* but not *NAT3* (Fig. 2*B*). These results indicate that Nt-acetylation is indeed important for Nma1 and Nma2 mediated NAD^+^ biosynthesis. Interestingly, NatA-specific Nmnat strains (*MSD, WT*) had lower basal levels of NAD^+^ compared to the NatB-specific Nmnat strains (*MD, WT*) (Fig. 2*B*). It is possible that NatA-mediated acetylation of Nmnats is less efficient compared to NatB. Potential NatA substrates include 25% of proteins vs 15% for NatB, possibly indicating NatA has a larger workload [51]. Additionally, most potential NatB substrates are indeed NatB substrates and are acetylated to near 100% efficiency, whereas NatA does not acetylate all of its potential substrates, and the level of acetylation on a NatA-specific substrate can also be different [28, 29].

We also addressed why NatB mutants have reduced levels of Nmnat. We previously showed that Nma1 and Nma2 proteins in NatB mutant cells were almost 100% non-Nt-acetylated [21], suggesting non-Nt-acetylated Nmnt proteins are more susceptible for protein degradation. Since the N-end rule pathways are closely associated with the degradation of Nt-acetylated peptides, and that Nmnat protein levels are correlated with NAD^+^ levels, we first examined whether deleting specific E3 ligases in the N-end rule pathways would restore NAD^+^ levels. Deleting known E3 ligases in N-rule pathways did not restore NAD^+^ levels (Fig. 3*A* and 3*B*). In addition, the degradation rates (half-live, t_1/2_) of Nma1 and Nma2 proteins were not significantly different when comparing non-Nt acetylated Nmnat (in NatB mutant) with Nt-acetylated Nmnat (in WT) (Fig. 3*C* to 3*H*). These results suggest that post-translational degradation by the N-rule pathways is not a major cause of Nmnat protein reduction in NatB mutants and that absence of Nt-acetylation did not significantly affect the half-lives of Nmnats. However, we cannot rule out the possibility that other E3 ligases or the ubiquitin-independent proteasomal degradation pathway may play a role. It has been show that the 20S proteasome (core subunit) can degrade ∼20% of all cellular proteins independent of ubiquitin ligation and that proteins with disordered regions are preferred 20S proteasome substrates [52, 53].

Our studies and previous reports suggest non-Nt-acetylated Nmnats may be more susceptible to co-translational degradation. Based on gel-filtration chromatography Nmnat proteins Nma1 and Nma2 likely function as tetramers [13–15]. While it was suggested that Nma1 likely functions as a homotetramer [13], Nma1 and Nma2 also co-immunoprecipitate suggesting other stoichiometry may exist [21, 54]. Furthermore, Nt-acetylation has been shown to be important for protein-protein interactions and can facilitate up to a 1000-fold increase in affinity [55, 56], which may be important for mediating co-translational complex formation and folding. It is therefore possible that NatB deletion mutants have low Nma1 and Nma2 protein level because co-translational complex formation is partially mediated through Nt-acetylation, and in the absence of Nt-acetylation the association of these nascent peptides with their partners (including chaperons) is decreased, leaving key residues vulnerable to degradation factors. Moreover, an intrinsically disordered region has been predicted at the N-terminus of Nma1 and Nma2 [45]. Intrinsically disordered regions of proteins lack stable tertiary structure. These regions are also functionally important for mediating protein-protein interactions and complex formation [45, 57]. It is possible that Nt-acetylation increases the hydrophobicity of the disordered region which assists the hydrophobic packing as seen with the formation of the E2-E3 complex Ubc12 and Dcn1 [58, 59]. Intrinsically disordered regions are also important features for mediating interactions with co-translational chaperones like Hsp70 [60]. Nascent Nma1 and Nma2 are substrates of Hsp70 and other chaperones [60, 61], and it is possible that Nt-acetylation promotes this interaction. In line with this, Nt-acetylation of Hsp90 subunits is required to mediate interactions with its substrates, and absence of Nt-acetylation increases degradation of both Hsp90 subunits and its substrates [33]. Weakened Nma1 and Nma2 interaction with chaperones is likely to increase misfolding of these proteins possibly leading to co-translational degradation. Interestingly, yeast and human Nmnats have a protective role against proteotoxicity, indicating Nmnats have a chaperone role as well [25, 27, 62]. Lastly, protein disorder also plays an important role in protein degradation by the proteasome. ATPase activity of the proteasome unfolds protein for translocation into the chamber. Translocation needs to be faster than refolding of the protein. In this sense disordered regions play an important role in mediating degradation because they promote an unfolded state that is efficiently translocated into proteasome for degradation [63].

Although Nt-acetylation has not been directly linked to the regulation of co-translational degradation, our studies of Nmnats suggest the status of Nt-acetylation may play a role. Co-translational degradation helps eliminate improperly folded translation intermediates whereas Nt-acetylation may facilitate proper folding and complex formation of newly synthesized polypeptides. It was shown that unassembled proteins are prone to degradation through exposure of degrons that are normally shielded when in complex [64]. Alternatively, studies in bacteria, yeast and mammalian cells suggest many multi-subunit complexes are built during translation, where proteins undergoing translation already begin interacting with their partners [65–67]. Preventing co-translational binding and folding often results in degradation of the translating partners, suggesting a competition between protein folding and complex formation with degradation. Nt-acetylation of Nma1 and Nma2 is likely important for mediating transient interactions or complex formation with proteins that either help protein folding or block key residues from being targeted by protein degradation machinery. This may help answer an interesting observation that in a normal cell Nma1, Nma2 and most other NatB substrates are always Nt-acetylated [21, 29]. It is likely that non-acetylated populations of these proteins exist but are hard to identify because they are degraded before reaching maturity.

There are 3 different Nmnat proteins in humans: Nmnat1, Nmnat2, and Nmnat3 [68]. Sequence conservation between human and yeast Nmnats ranges from 39 to 47% for Nma1 and 36 to 42% for Nma2. Yeast Nmnats have a ∼150 amino acid N-terminal extension not seen with human Nmnats [68]. Based on this information it is possible that Nt-acetylation of Nmnats in human does not fulfill the same purpose. However, even with the large difference in sequences, Nmnats from yeast and human have two functions that are well conserved. The first function is that they wield dual NAMN- and NMN-adenylyltransferase activities. Secondly, both yeast and human Nmnats have a chaperone role mediating protein folding [25, 27, 62]. Based on the protein sequences, Nmnat1 is the only human Nmnat that is likely a potential NatB substrate. Nmnat2 and Nmnat3 are potential substrates of NatA and NatX, respectively. It would be interesting to know if Nt-acetylation plays an important role in regulating human Nmnats as well. In conclusion, our studies help understand the mechanisms of NatB mediated regulation of NAD^+^ biosynthesis by modulating Nt-acetylation of Nmnats. It would be informative to further investigate whether and how Nt-acetylation of Nmnat affects its complex formation and function. These studies may also provide insight into the regulation of NAD^+^ homeostasis and the molecular basis of disorders associated with aberrant NAD^+^ metabolism.

## EXPERIMENTAL PROCEDURES

### Yeast Strains, Growth Media and Plasmids

Yeast strain BY4742 *MATα his3Δ1 leu2Δ0 lys2Δ0 ura3Δ0* acquired from Open Biosystems [69] was used for this study. Standard media such as synthetic complete (SC) media, synthetic minimal media (SD), and yeast extract-peptone-dextrose (YPD) rich media were made as described [70]. NA/NAM-free SD and NA/NAM-free SC were made by using niacin-free yeast nitrogen base (Sunrise Science Products). SC was used for most experiments unless noted. Gene deletions were generated by replacing the coding regions of wild type (WT) genes with gene-specific PCR products generated using either the pAG32-hphMX4 (hygromycin resistance) or reusable loxP-*kan^r^*-loxP cassettes [71, 72]. Multiple deletions were carried out by removing the *kan^r^* marker using a galactose-inducible Cre recombinase [71]. *NMA1*, and *NMA2* were tagged by the HA tag directly in the genome using the pFA6a-3HA-HIS3MX system as previously described [73, 74]. Plasmids carrying *NMA1* and *NMA2* were generated by ligating *NMA1* (-803 to +1496) and *NMA2* (-522 to +1702) PCR fragments, which include promoter and terminator regions, into pRG205 and pRG207 (cut with KpnI and HindIII enzymes), and the plasmids were linearized with AscI and integrated into the genome as described [75]. Using these parent plasmids, mutation and insertion variants of *NMA1* and *NMA2,* NMA1-MSD (M_1_D_2_ ➔ M_1_S_2_D_3_, Ser inserted between the first Met and the second Asp), NMA2-MSD (M_1_D_2_➔ M_1_S_2_D_3_), and *NMA1*-int HA-tag (K_30_S_31_ ➔ K_30-YPYDVPDYA-_S_40_; HA tag added between K_30_ and S_31)_ were generated using the NEB Q5^®^ Site-Directed Mutagenesis Kit. Strains carrying Not4-AID*-9myc was generated by integration using PCR product amplified from the pKan-AID*-9myc plasmid [44]. Plasmid pRG207-p*ADH1*-*OsTIR1-9MYC* was made by ligation of the p*ADH1-OsTIR1-9MYC* fragment from pNHK53 into pRG207 [43]. This allows integration of pRG207-p*ADH1*-*osTIR1-9MYC* to the *LYS2* locus since our strain does not have the *URA3* locus required for pNHK53 integration. The pKan-AID*-9myc and pNHK53 plasmids were obtained from the National BioResource Project–Yeast. The pRG205 and pRG207 plasmids were obtained from Addgene. All plasmid constructs were verified by DNA sequencing.

### NAD^+^ and NADH measurements

NAD^+^(H) was measured by enzymatic cycling assay as previously described [76, 77]. In brief, cells were grown to early-mid log phase (*A*_600_ of ∼1), and 1 *A*_600_ unit (1 *A*_600_ unit = 1 x10^7^ cells/ml) of cells was collected in duplicates. Acid extraction was performed in one tube to obtain NAD^+^, and alkali extraction was performed in the other to obtain NADH for 40 minutes at 60°C. Amplification of NAD^+^ or NADH in the form of malate was carried out using 3 µl or 4 µl of neutralized acid or alkali extracted lysate in 100 µl of cycling reaction for 1 hour at room temperature, and the reaction was terminated by heating at 100°C for 5 minutes. Once cooled, malate produced from the cycling reaction was converted to oxaloacetate and then to aspartate and α-ketoglutarate by the addition of 1 ml malate indicator reagent for 20 minutes at room temperature. The reaction produced a corresponding amount of NADH as readout, which was measured fluorometrically with excitation at 365 nm and emission monitored at 460 nm. Standard curves for determining NAD^+^ and NADH concentrations were obtained as follows: NAD^+^ and NADH were added into the acid and alkali buffer to a final concentration of 0, 2.5, 5, and 7.5 µM, which were then treated with the same procedure along with other samples. The fluorometer was calibrated each time before use with 0, 5, 10, 20, 30, and 40 µM NADH to ensure that the detection was within a linear range.

### RNA extraction, cDNA synthesis and quantitative PCR (qPCR)

Cells were grown in SC to early-mid log phase (*A*_600_ of ∼1) and 40 *A*_600_ units of cells were collected. Total RNA was isolated using GeneJET RNA purification Kit (Thermo Scientific) and cDNA was synthesized using QuantiTect Reverse Transcription kit (Qiagen) according to the manufacturer’s instructions. For each qPCR reaction, 100 ng of cDNA and 500 nM of each primer were used. qPCR reaction was run on Roche LightCycler 480 using LightCycler 480 SYBR green I Master Mix (Roche) as previously described [19]. The target mRNA transcripts were normalized to *TAF10* transcript levels [78].

### Cycloheximide chase and Western blots

For cycloheximide chase studies, cells were grown in SC for 5-6 hrs to early-mid log phase (*A*_600_ of ∼1) before either harvesting (0 hour time point) or treating with 0.4 mg/ml cycloheximide (CHX) followed by harvesting at 1 and 2 hours later. This step was skipped if measuring basal protein level. Cell lysates for Western blot were made as described [33]. Briefly, 1 *A*_600_ unit of c ells was pelleted and supernatant was removed. Treatment with 800 µl of 2 M Lithium acetate was carried out on ice for 5 minutes, followed by pelleting and the removal of supernatant. 800 µl of 0.4 M NaOH was then added for 5 minutes on ice, followed by pelleting and removal of supernatant. Cells were lysed by boiling the pellet in 80 µl HU buffer (8 M urea, 0.2 M Tris-HCl pH 6.8, 5% SDS, 1 mM EDTA, 0.1 M DTT, 0.005% bromophenol blue) with Roche EDTA free cOmplete Mini protease inhibitor cocktail at 70°C for 10 minutes. Lysates were clarified by centrifugation for 10 minutes and stored or loaded into a 10% SDS-PAGE. Equal volume (∼20 µl) of lysate was loaded in each lane. Following electrophoresis, protein was transferred to low fluorescence PVDF (GE). Blocking was carried out using OneBlockTM Western-FL Blocking Buffer (20-314), followed by blotting using a combination of anti-HA Rabbit (Cell Signaling 3724S), anti-c-Myc Rabbit (Novus NBP2-52636) and anti-PGK1 Mouse (Invitrogen 459250). Secondary antibodies Goat anti-Rabbit IgG Alexa Fluor Plus 647 (A32733) and Goat anti-Mouse IgG Alexa Fluor 555 (A32727) were used to visualize the protein using an Amersham Imager 600RGB imaging system (GE). Protein intensity was calculated using the Amersham Imager 600 software.

### NA/NAM cross-feeding spot assays

The *bna6Δnrk1Δnrt1Δ* mutant (which could utilize NA and NAM) was used as “recipient cells” and yeast strains of interest were used as “feeder cells”. First, recipient cells were plated onto NA/NAM free SD as a lawn (∼10^4^ cells/cm^2^) as previously described [21]. Next, 2 µl of each feeder cell strain (∼10^4^ cells, cell suspension was made in sterile water at *A*_600_ of 1) was spotted onto the lawn of recipient cells. Plates were then incubated at 30°C for 3 days. Extent of the recipient cell growth indicates the levels of NA/NAM released by feeder cells.

### Polysome profiling and isolation of mRNA

Polysome profiling was carried out using early-mid log phase cells grown in 100 ml SC or YPD as described [47–49] with modifications. Once cells reached approximately *A*_600_ of 1, 0.2 mg/ml cycloheximide was added to the culture and allowed to grow for 10 minutes. Cells were collected by centrifugation and washed once in polysome extraction buffer (20 mM Tris-HCl pH 7.5, 50 mM KCl, 10 mM MgCl_2_, 5 mM DTT, and 0.2 mg/ml cycloheximide). Cell pellets were flash frozen in liquid nitrogen and stored at -80°C overnight. 10%-50% sucrose gradients were made as followed: ∼2.2 ml of 10%, 20%, 30%, 40%, and 50% sucrose solutions (20 mM Tris-HCl pH 7.5, 100 mM NaCl, 5 mM MgCl_2_, 0.1 mg/ml cycloheximide, 1mM DTT and 5 U/ml RiboLock RNase Inhibitor (EO0381)) were overlaid with freezing of each layer at -80 °C before the next was added. Once all layers were added the gradients were kept indefinitely at -80 °C. 24 hours prior to use, gradients were thawed at 4°C to become linear. Next day, pre-chilled 500 µl glass beads and 400 µl polysome extraction buffer with the addition of 1X Roche EDTA-free cOmplete Mini protease inhibitor cocktail and 1 U/µl RiboLock RNase Inhibitor was added to the cell pellet. Beads beating was carried out continuously at 3,000 RPM for 15 minutes at 4°C. The supernatant was transferred to a new tube and clarified for 20 minutes at 21,130x g at 4°C. About 400 µl cell lysate was layered onto the gradients and centrifugated at 45,000 RPM in a SW41Ti swinging bucket rotor for 2.5 hours. Sucrose gradients were collected using a Brandel Density Gradient Fractionation System into 10 fractions (∼1 ml for each fraction). RNAs in each fraction were concentrated by addition of 600 µl TRIzol and 240 µl chloroform to 600 µl from each fraction. Samples were mixed by vortexing and aqueous and organic layers were separated by centrifugation at 21,310x g for 15 minutes at 4°C. 500 µl of the upper layer was transferred to a new tube containing 1 ml isopropanol and 2 µl of 15 mg/ml glycoblue. RNA was precipitated overnight at -20°C and pelleted by centrifugation at 21,310x g for 15 minutes at 4°C. Supernatant was removed and the pellet was washed once with 1 ml ice-cold 70% ethanol followed by pelleting and removal of the supernatant. Pellet was air dried for 5-10 minutes at room temperature and dissolved in 12 µl of nuclease-free water. cDNA synthesis and qPCR was carried out as described above with the following modifications. For each fraction, 1 µg of mRNA was used for cDNA synthesis in every 20 µl reaction. 2 µl of the resultant cDNA (∼100 ng) was used for each qPCR reaction. The amount of gene-specific mRNA in each fraction was calculated as described [48, 50]. In brief, relative levels of gene-specific cDNA in each fraction were first normalized to fraction 1 and then normalized to total amount (the sum of 10 fractions) to obtain the proportion (%) of the target gene found in each fraction. As a result, relative abundance and distribution of specific mRNA in all 10 fractions were shown for each gene.

## Acknowledgments

We thank Dr. C. Fraser for assistance with polysome profile studies and Dr. J. Roth and Dr. S. Collins for suggestions and discussions. This study was supported by the National Institutes of Health, NIGMS (GM102297).

## Author contributions

T. Croft and S. -J. L. conceptualization; T. Croft, P. V., C. J. T. R., B. G., T. Cater, and S. -J. L. data curation; T. Croft and S. -J. L. formal analysis; T. Croft, P. V., C. J. T. R., B. G., and T. Cater validation; T. Croft, P. V., C. J. T. R., B. G., and T. Cater investigation; T. Croft, P. V., C. J. T. R., and S. -J. L. visualization; T. Croft, P. V., C. J. T. R. and S. -J. L. methodology; T. Croft and S. -J. L. writing-original draft; T. Croft, P. V., C. J. T. R., B. G., and S. -J. L. writing-review and editing; S. -J. L. supervision; S.-J. L. funding acquisition; S.-J. L. project administration.

